# Spatial-temporal clustering analysis of yaws on Lihir Island, Papua New Guinea to enhance planning and implementation of eradication programs

**DOI:** 10.1101/352542

**Authors:** Eric Q. Mooring, Oriol Mitjà, Megan B. Murray

## Abstract

**Background:** In the global program for the eradication of yaws, assessments of the prevalence of the disease are used to decide where to initiate mass treatment. However, the smallest administrative unit which should be used as the basis for making decisions is not clear. We investigated spatial and temporal clustering of yaws to help inform the choice of implementation unit.

**Methodology/Principal findings:** We analyzed 11 years of passive surveillance data on incident yaws cases (n = 1448) from Lihir Island, Papua New Guinea. After adjusting for age, sex, and trends in health-seeking, we detected three non-overlapping spatiotemporal clusters (p < 1 × 10^−17^, p = 1.4 × 10^−14^, p = 1.4 × 10^−8^). These lasted from 28 to 47 months in duration and each encompassed between 4 and 6 villages. We also assessed spatial clustering of prevalent yaws cases (n = 532) that had been detected in 7 biannual active case finding surveys beginning in 2013. We identified 1 statistically significant cluster in each survey. We considered the possibility that schools that serve multiple villages might be loci of transmission, but we found no evidence that incident cases of yaws among 8- to 14-year-olds clustered within primary school attendance areas (p = 0.684).

**Conclusions/Significance:** These clusters likely reflect transmission of yaws across village boundaries; villages may be epidemiologically linked to a degree such that mass drug administration may be more effectively implemented at a spatial scale larger than the individual village.

**Author Summary:** The World Health Organization aims to eradicate yaws using mass drug administration (MDA), which consists of treating everyone in an administrative unit with antibiotics. The administrative unit in a country which is used as the basis for making decisions about implementing MDA is called the implementation unit. Prevalence assessments are used to identify endemic communities for mass treatment programs, but the spatial scale (e.g. village, sub-district, district, or province) at which mass treatment should be implemented is currently unclear. The choice of implementation unit depends on many factors; one of these is the underlying transmission patterns of the disease. Using data from Lihir Island, Papua New Guinea, we found that geographic clusters of yaws often spanned multiple villages. These clusters likely reflect transmission of the infectious disease across village boundaries and suggest that it may be best to implement MDA at a spatial scale larger than the individual village, for example at sub-district level.

## Introduction

Yaws, a bacterial disease caused by *Treponema pallidum* subspecies *pertenue*, causes skin lesions and arthralgia, most commonly in school-age children [1]. Yaws spreads via direct skin-to-skin contact. While yaws was widespread in tropical areas in the first half of the twentieth century, the disease currently persists primarily in Melanesia and parts of west and central Africa [2]. The discovery that oral azithromycin treats yaws as effectively as injectable penicillin sparked renewed interest in eradicating this neglected tropical disease [3,4]. Yaws often enters a latent stage during which patients are no longer infectious or symptomatic, but still infected and at risk of the disease reactivating and once again being infectious. Eradicating yaws necessarily requires treating all infections, including latent ones. The World Health Organization (WHO) yaws eradication strategy calls for 1 or more rounds of mass drug administration (MDA) in all yaws-endemic areas; that is, treating with azithromycin nearly everyone in these areas, regardless of individuals’ disease status. MDA aims to cure yaws even in latently infected individuals, thereby preventing them from later becoming infectious.

Before implementing yaws elimination programs, public health officials need to decide the spatial scale at which to conduct MDA. They refer to the geographic level (e.g. province, district, sub-district or village) with respect to which they decide to start and stop MDA as the “implementation unit” [5]. The WHO initially recommended that the village or community should be the implementation unit [4]. This recommendation was informed, in part, by research concluding that village-level yaws prevalence is a stronger risk factor for yaws than household-level prevalence [6,7]. A more recent review argued that conducting yaws prevalence surveys at the village level is impractical and that the implementation unit for the initial round of MDA should instead be a unit with a population of 100,000 to 250,000 people and that for subsequent treatment rounds the implementation unit could be lowered to the village [8]. The choice of implementation unit is an unresolved challenge in formulating yaws eradication strategy and reflects a lack of research on the spatial epidemiology of yaws. Understanding yaws transmission and clustering not only helps inform the choice of implementation unit, but also is needed to design statistically rigorous assessments of yaws prevalence within each implementation unit. Because clustering of an infectious disease can occur at multiple spatial scales simultaneously [9], research is needed on yaws spatial epidemiology at multiple scales. In this study we assess whether clusters (i.e. areas of disproportionately high prevalence or incidence) of yaws extend across neighboring villages. Spatial-temporal clusters of disease extending across villages may indicate that appreciable levels of disease transmission occur between villages. While most yaws studies have relied on cross-sectional data, we leverage more than a decade of clinical records to include a temporal component in our analysis. Finally, given that yaws is predominantly a disease of school children and might be expected to be spread at schools, we investigate whether primary school attendance areas help explain the spatial distribution of yaws.

## Methods

### Study setting

We conducted this study on Lihir Island, a 200 km^2^ tropical island in New Ireland Province, Papua New Guinea. Fig 1 shows that the island’s mountainous interior is largely uninhabited and the population of approximately 12 500 Lihirians lives in coastal villages linked by a road that encircles the island. Even the most geographically distant pair of villages are only 35.2 km apart by road. Approximately 4680 migrants live on the island. The island houses a large-scale gold mine. A network of government-run aid posts, a clinic at a Catholic mission station, and the Lihir Medical Centre (LMC) (a hospital that serves both mine personnel and the public) deliver health care services. Yaws has long been endemic in the region; Dr. Robert Koch visited Lihir in 1900 and subsequently reported that he observed yaws throughout the Bismarck Archipelago, the group of islands that comprise the northeastern part of present-day Papua New Guinea [10,11].

**Fig 1.**
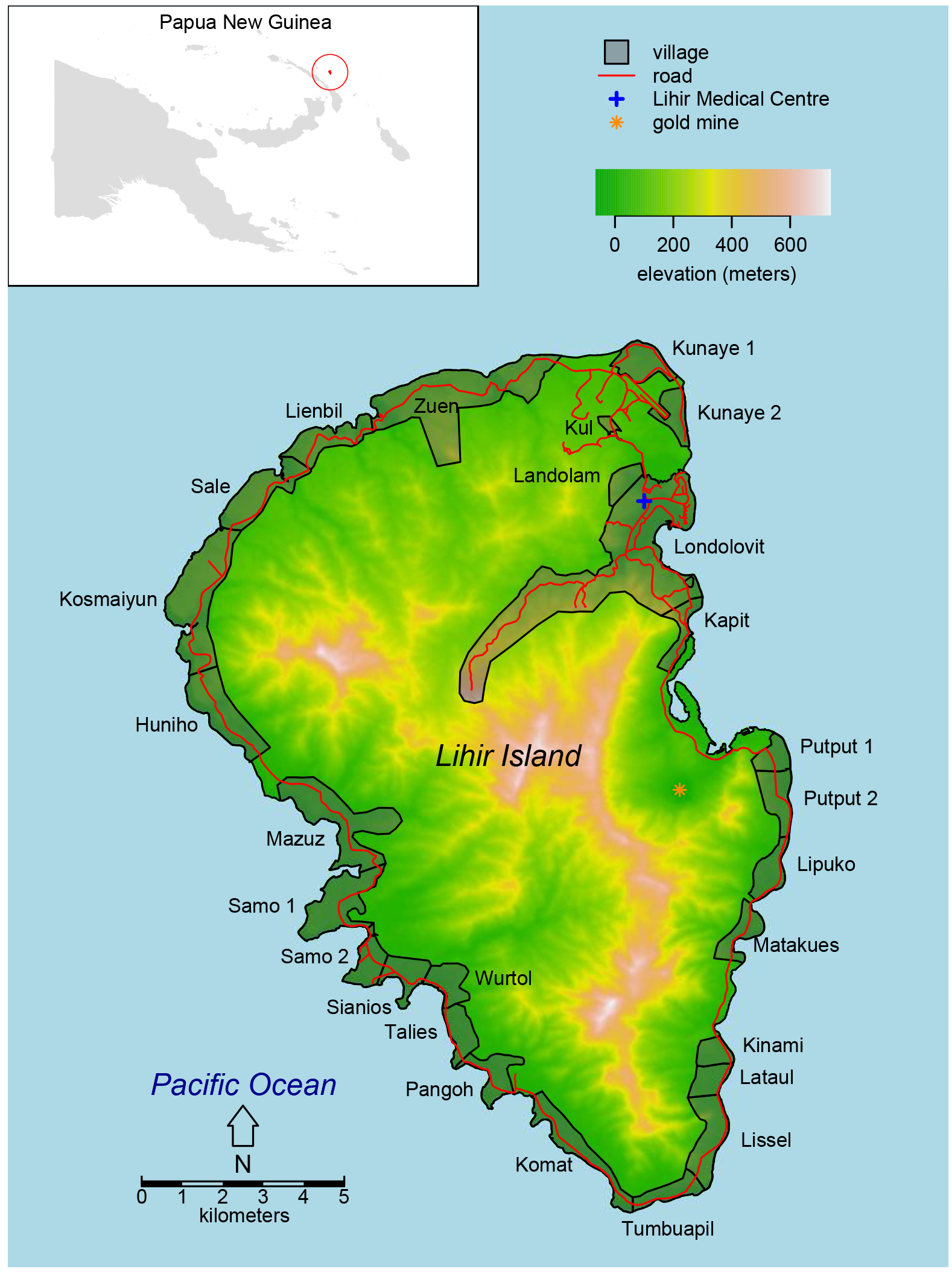
Map of Lihir Island. The top right inset shows the location of Lihir in Papua New Guinea. Note that Landolam is technically not a village but rather an informal migrant settlement. Elevation data was obtained from the Shuttle Radar Topography Mission [12]. The Papua New Guinea map was obtained from GADM database [13].

### Spatial-temporal clusters of incident yaws from outpatient records

We queried the LMC electronic medical record to identify all outpatients who had been diagnosed with yaws between early April 2005 and 29 May 2016. We included only those patients whose reported place of residence was a Lihir village. LMC typically recorded migrants to Lihir (even those living there indefinitely) as being from their place of origin, not the Lihirian village where they currently live.

To be considered an incident yaws case at the LMC, patients must present with a skin lesion consistent with yaws and have positive results on both the *Treponema pallidum* hemagglutination assay (TPHA) and the rapid plasma reagin (RPR) test. The THPA is a highly specific biomarker that provides definitive diagnosis of prior or current treponemal infection, while RPR, despite being less specific, indicates current disease activity [14].

We defined a cluster as a set of 1 or more villages where we observed more incident yaws cases in a time period than expected, given the spatial and temporal distribution of all outpatient visits to the LMC by Lihir village residents and the age and sex distribution of these patients. To detect statistically significant non-spatially overlapping clusters of outpatient yaws diagnoses, we conducted a retrospective space-time analysis in SaTScan version 9.4.4 using the discrete Poisson model [15–17]. We defined the spatial relationships between villages by the road distances between village centroids. Following standard practice for using SaTScan, we set the maximum allowed cluster duration to be half the duration of the study period. Similarly, we constrained the maximum spatial size of a cluster to encompass villages accounting for no more than half the outpatient visits over the course of the study period.

### Spatial clusters of prevalent yaws from active surveys

Starting in April 2013, we implemented a round of mass drug administration with azithromycin on Lihir to demonstrate the feasibility of eliminating yaws [18]. All people older than 2 months were offered treatment regardless of symptoms. At the same we administered azithromycin, we examined the participants to identify all suspected yaws cases. We subsequently screened the population every 6 months to identify and treat symptomatic yaws cases and their contacts. In this study, we analyze data through the seventh survey of active case finding, which took place in April and May 2016. We collected venous blood samples from consenting individuals suspected of having yaws and performed TPHA and RPR testing on these samples. We swabbed the ulcers of all suspected yaws cases in surveys 3 through 7. Polymerase chain reaction (PCR) tests were performed on these swabs. The test consists of amplifying three *T. pallidum* gene targets: *tp0548*, tpN47 (*tp0574*), and a *pertenue*-specific region of the tprL (*tp1031*) gene [19].

In contrast to individuals in the outpatient data from the LMC, we categorized screened individuals in the active case finding data by their current village of residence, without regard to migrant status. We also included as a village in this analysis an informal migrant settlement. We excluded from the analysis cases in villages during surveys where the number of individuals screened was missing for that village.

To assess spatial clusters based on data from active case finding with serological confirmation, we analyzed data from each survey separately using the spatial-only discrete Poisson model in SaTScan. Clusters were villages or groups of villages where the proportion of screened individuals who had yaws was greater than expected. We constrained the maximum cluster size such that no cluster contains more than half of the screened population.

### Clustering of incident yaws cases within primary school attendance areas

Using data on the village of residence of students at each Lihir primary school in 2013, we defined primary school attendance areas based on the most frequent primary school affiliation of primary school-enrolled children from each village.

In this analysis, we restricted the outpatient visit data to 8-to 14-year-olds. We calculated for each village the number of incident yaws cases divided by the total number of outpatient visits. Next, we calculated the F-statistic from weighted analysis of variance (ANOVA) to quantify the degree to which primary school attendance areas account for between-village differences in the number of yaws cases per outpatient visit. We weighted each proportion by the inverse of its variance. Because the inverse of the variance is not defined when there are no yaws cases among 8- to 14-year-olds from a given village in a given year, we added 2 yaws cases and 4 outpatient visits to every village only when calculating weights [20].

Finally, to test whether clustering of yaws cases in primary school attendance areas deviated from the null hypothesis (that clustered cases in villages served by a given primary school result from their spatial clustering rather than attendance at the same school), we permuted the primary school attendance area assignments while maintaining the condition that each primary school serves a sequential set of villages along the circumference of the island and calculated the F-statistic for each permutation using weighted ANOVA, thereby constructing an empirical distribution from which we calculated p-values [21]. Analyses were conducted in R version 3.4.2 [22].

### Ethics approval and informed consent

The protocol was approved by the National Medical Research Advisory Committee of the Papua New Guinea National Department of Health (MRAC no. 12.36).

## Results

### Passive case detection and spatial-temporal clusters of incident yaws cases

From 11 years of routinely collected clinical data on residents of Lihir villages, we identified 2365 distinct symptomatic yaws cases but excluded 917. Our main analysis is based on 1448 cases of yaws among 1271 patients (Fig 2A). During this period, Lihirians made a total of 288 729 outpatient visits to the LMC (Fig 2B; S1 Fig). The median age at diagnosis of the yaws cases was 9 years (interquartile range: 6 – 13). The cumulative number of yaws patients per village varied widely, ranging from 4 yaws cases out of 1390 outpatient visits from Huniho and 4 out of 3156 from Lienbil to 234 out of 32859 from Kunaye 1.

**Fig 2.**
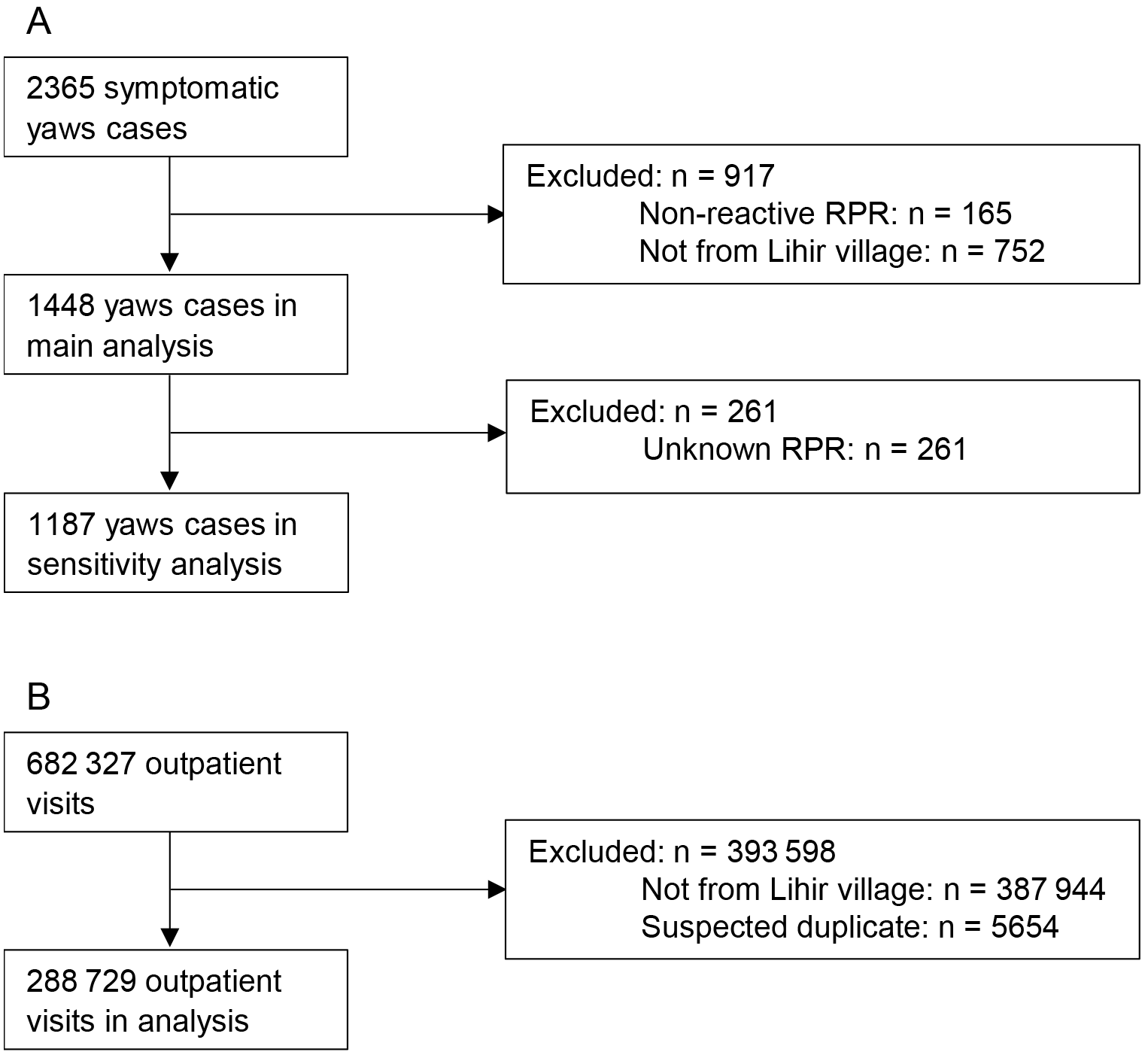
Flow diagrams for data in analysis of passively detected yaws. (A) Exclusions of outpatient passively detected yaws cases and (B) exclusions of all outpatient visits at Lihir Medical Centre.

We detected 3 statistically significant spatial-temporal clusters (Fig 3, S2 Fig). The most statistically significant cluster (p < 1 × 10^−17^) lasted from July 2009 into March 2012 and encompassed 6 villages at the southern tip of Lihir (cluster 1); the most distant pair of villages in that cluster are 10.7 km apart by road. The next most statistically significant cluster (p = 1.4 × 10^−14^) lasted from the beginning of the study period (April 2005) until the beginning of March 2009 and occurred in 4 villages located along the east coast of Lihir (cluster 2) and spanned over a road distance of 5.1 km. Finally, the third statistically significant cluster (p = 1.4 × 10^−8^) lasted from January 2014 through the end of the study period in May 2016. It comprised 4 villages located in the northeast corner of Lihir (cluster 3) which are at most 7.4 kilometers apart by road. Our analysis of spatial-temporal clustering incident yaws cases was robust to a range of assumptions, including restricting only to yaws cases for which a positive RPR result was recorded in the medical record (S1 Table), not adjusting for age and sex (S2 Table), and using SaTScan’s case-only space-time permutation method instead of the discrete Poisson method [23] (S3 Table and S4 Table). While the exact timing and duration of clusters varied somewhat between the different analyses, all found evidence of 3 high statistically significant spatial-temporal clusters, and the temporal order and general locations of those clusters were unchanged.

**Fig 3.**
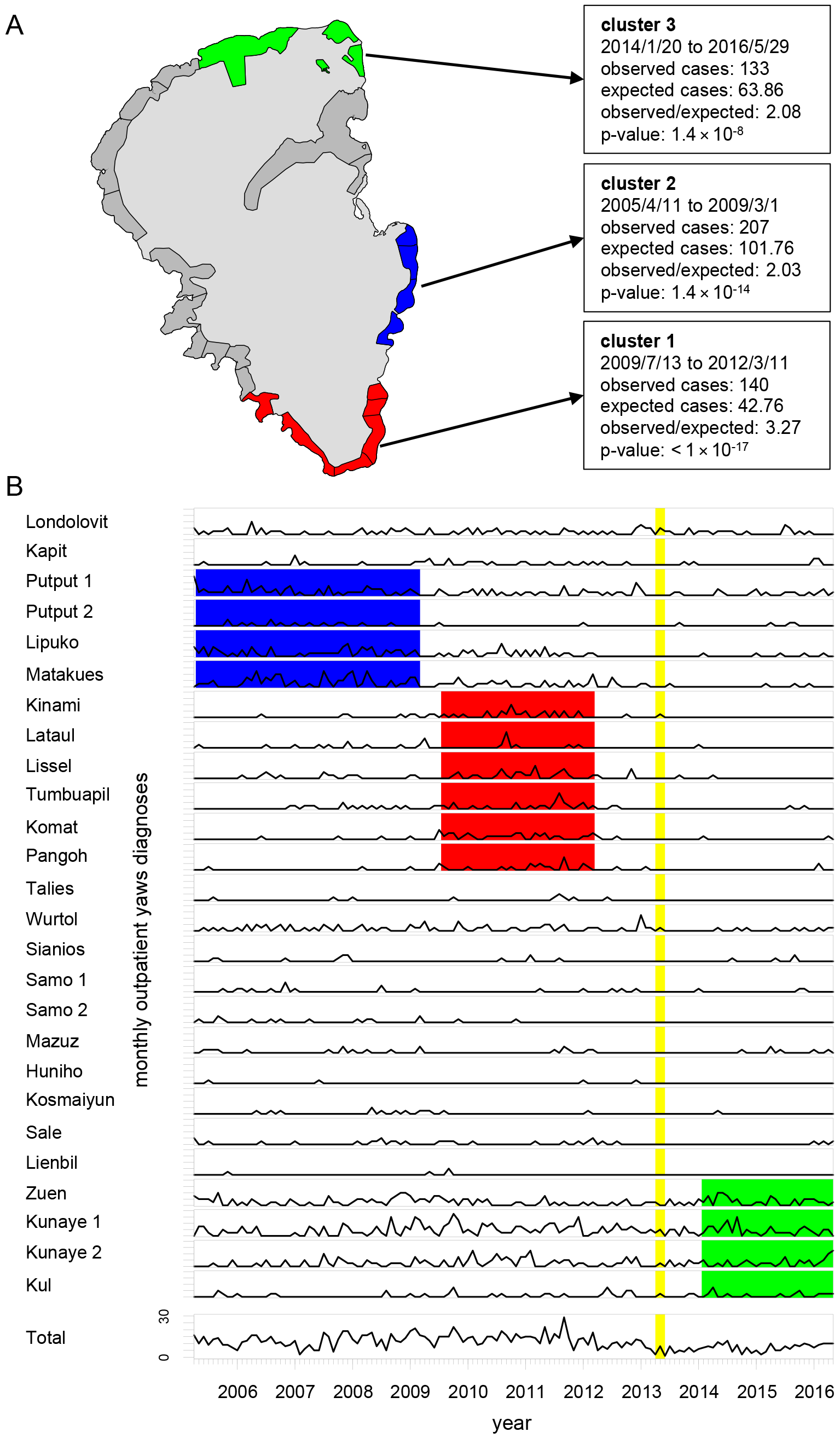
Spatial-temporal clusters of passively detected yaws cases. (A) Map of results of spatial-temporal Poisson SaTScan analysis of serologically confirmed incident yaws cases, adjusted for age and sex. (B) Times series of outpatient yaws cases aggregated by month by village and for all villages combined. The y-axes for the village-specific time series range from 0 to 8. The villages are ordered to match their sequential order around the circumference of Lihir. Red, blue, and green rectangles correspond to spatial-temporal clusters 1, 2, and 3, respectively. The vertical yellow bar corresponds to when mass drug administration was implemented on Lihir.

### Active case finding and spatial clustering of prevalent yaws cases

Out of 95 353 screenings during the 7 biannual active case finding surveys (mean of 13 622 individuals screened per survey), we identified 568 serologically confirmed yaws cases, of which 36 were excluded (Fig 4). More than half of all serologically confirmed cases were identified during the first (pre-MDA) round of screening. Before exclusions, the median age of the serologically confirmed cases was 12 years (interquartile range: 8 – 14). We detected 1 statistically significant cluster in each survey, at the α = 0.05 level of significance (Fig 5). The number of villages in each cluster varied widely from 1 to 16.

**Fig 4.**
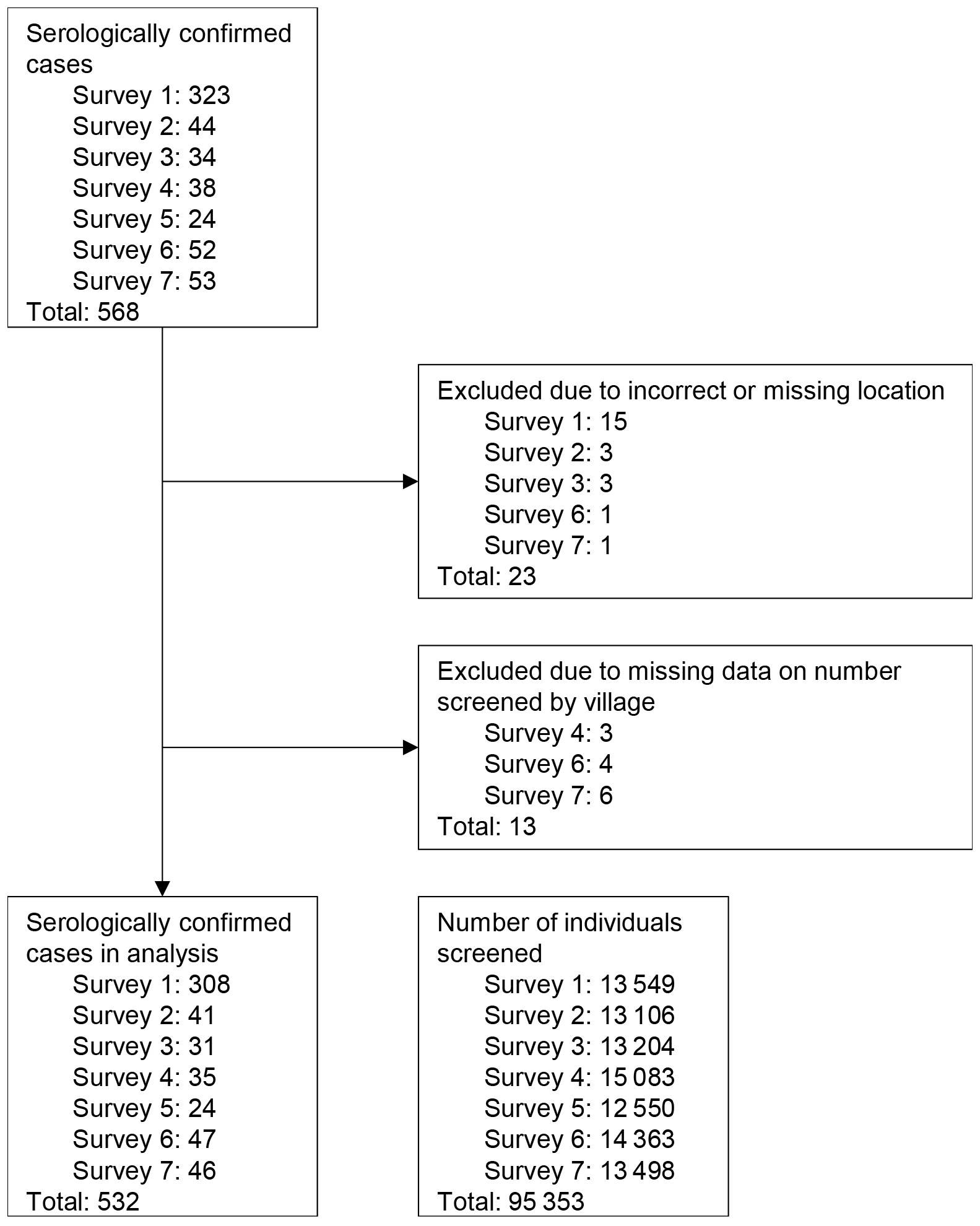
Flow diagram for data in analysis of prevalent yaws cases. The diagram lists serologically confirmed cases, excluded cases, and number of individuals screened by active case finding survey.

**Fig 5.**
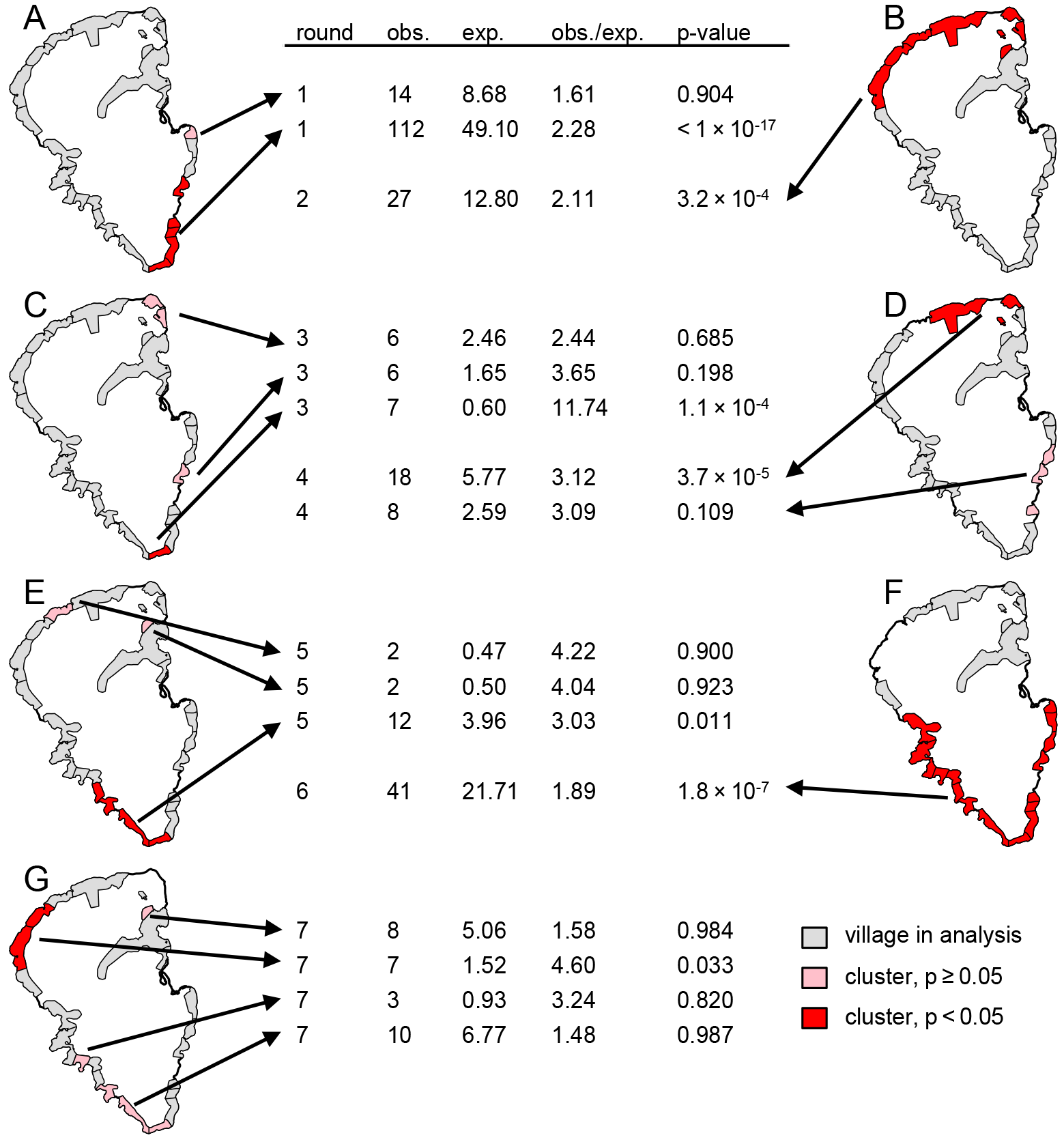
Spatial clusters of actively detected serologically confirmed yaws cases. Results of spatial-only discrete Poisson SaTScan analysis for serologically confirmed prevalent yaws cases identified via active case finding for each survey 1 through 7 (A ‒ G, respectively). Villages that are part of statistically significant spatial clusters are shaded in red and not statistically significant clusters are shaded in pink. Villages are shaded gray if they were part of the analysis in the corresponding survey (but not identified as part of a cluster). Villages are excluded from the map for each survey where the number of individuals screened in that survey in that village is unknown. The arrows point from each cluster to details describing the number of observed yaws cases in that cluster, the number of expected yaws cases, the ratio of observed to expected, and p-value for that cluster.

In a supporting analysis we repeated our active-case finding analysis using PCR-confirmed cases only. S3 Fig lists 116 PCR-confirmed cases and 11 exclusions by survey, and results for PCR-confirmed cases are shown in S4 Fig. The results from PCR-confirmed and serologically confirmed yaws cases are broadly similar in that most clusters of serologically-confirmed cases overlapped with a cluster of PCR-confirmed cases, but fewer of the clusters detected in the PCR-based analysis were statistically significant. Unlike in the analysis of serologically confirmed cases, there were no statistically significant clusters of PCR-confirmed yaws in the third or fifth surveys.

### Primary school attendance areas and yaws

We categorized each village into 1 of 7 primary school attendance areas. Four of the primary school attendance areas consisted of 4 villages, while the remaining 3 areas consisted of 1, 3, and 6 villages apiece (Fig 6A). We identified a total of 643 incident cases of serologically confirmed yaws among 8- to 14-year-olds, out of 32 202 visits by outpatients age 8 to 14. We did not find evidence for clustering of incident yaws cases among 8- to 14-year-olds within primary school attendance areas in individual years 2006 through 2015 (p-values ranged from 0.0687 in 2006 to 0.9881 in 2009), nor in the combined data (p = 0.6846) (Fig 6 and S5 Table).

**Fig 6.**
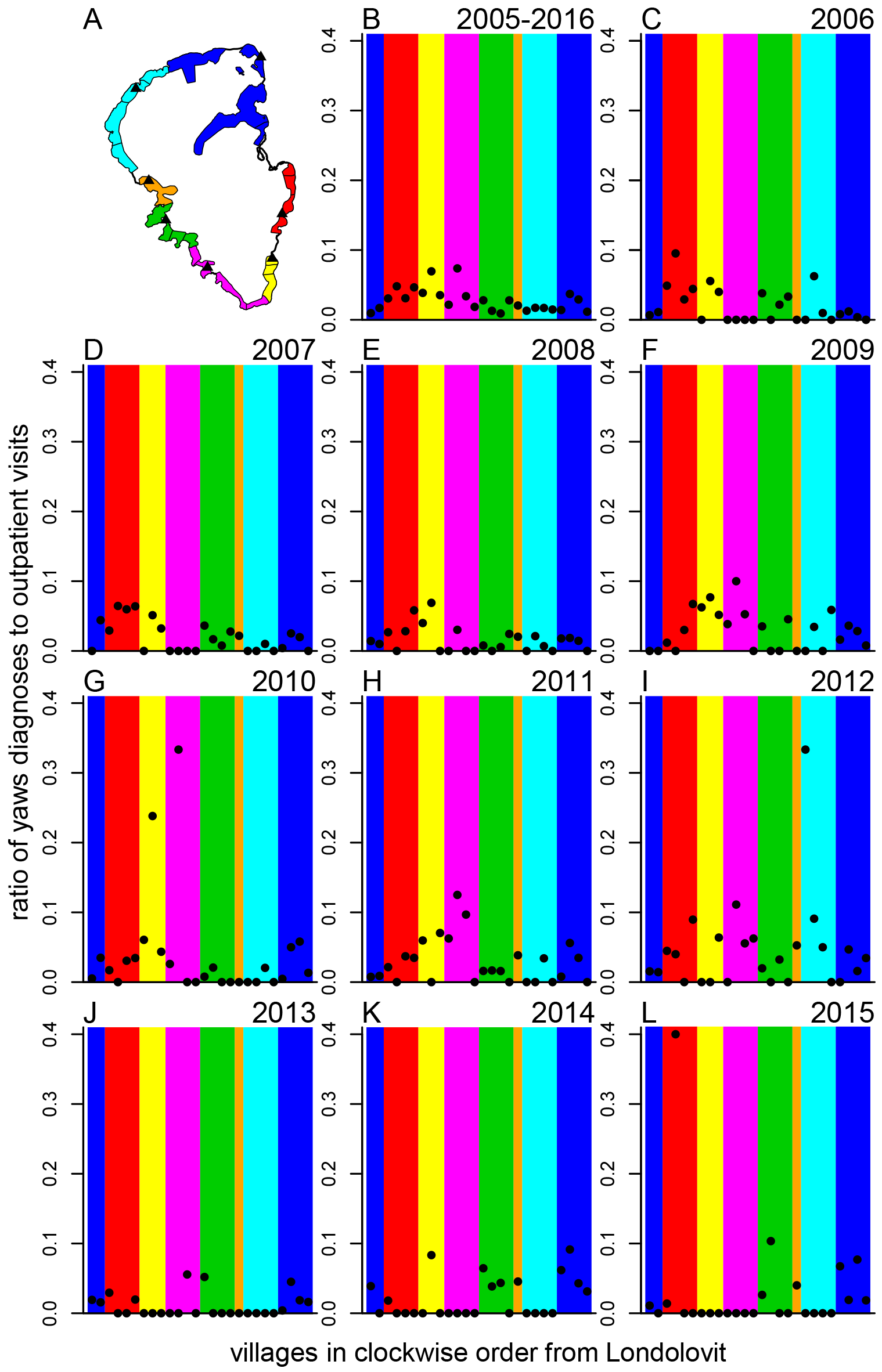
Passively detected yaws cases by primary school attendance area and year. (A) Map of Lihir villages with the colors corresponding to empirically derived primary school attendance areas. Each solid black triangle marks the location of a primary school. (B ‒ L) Yaws diagnoses among 8- to 14-year-olds as a proportion of outpatient visits for the entire study period (B) and each year 2006 to 2015 (C ‒ L, respectively). Each data point corresponds to a village and the background colors in each plot correspond to the primary school attendance areas illustrated in (A). The data points ordered left-to-right correspond to the clockwise sequences of villages around Lihir arbitrarily starting from Londolovit, the large village in the northeast quadrant of the island.

## Discussion

We found highly statistically significant spatial-temporal clusters of incident yaws cases, which stretched over multiple years and multiple villages. These clusters likely reflect transmission of yaws across village boundaries. The overall incidence of yaws can be viewed as the sum of an endemic and an epidemic component. The transmission dynamics of yaws on Lihir may consist of repeated and overlapping “outbreaks” of yaws that, when aggregated at the scale of the entire island, create the appearance of a persistently high-level endemic disease. The results of our analysis of purely spatial clusters (based on data from biannual surveys of active case finding) correspond in some respects to the results of our analysis of spatial-temporal clusters. For example, in the period January 2014 to May 2016 we identified a spatial-temporal cluster in northeastern Lihir that coincided with purely spatial clusters identified in May and October 2014 in the same area. More generally, our analysis of active case finding data found multi-village clusters of yaws in nearly all 7 surveys. This provides further evidence that at least some of the mechanisms that govern the spatial and temporal distribution of yaws operate at a scale larger than the village level. We did not find evidence for clustering of yaws within primary school attendance areas. While yaws transmission could be rare in an absolute sense within primary schools, a plausible alternative is that outside-of-school transmission simply overwhelms any signal from within-school transmission.

Our study has implications for yaws elimination: If spatial-temporal clusters are an inherent part of the transmission dynamics of yaws, then this suggests that even if interventions are implemented consistently, the path to elimination is unlikely to be smooth and clusters of yaws may still occur. If the background prevalence of yaws is lower, presumably any clusters would be more apparent. Also, our analysis suggests that the persistence of yaws following MDA cannot be attributed purely to a lack of control in a single location nor to reintroduction of yaws at just 1 location. Indeed, our findings are compatible with the fact that many of the cases of yaws that occurred following MDA were in people who were not treated during MDA [24]. Finally, if the observed clustering is due to appreciable inter-village yaws transmission, we suggest that villages may be epidemiologically linked to such a degree that they may not effectively serve as implementation units. If the recommended implementation unit size were 20 000 to 50 000 people, then Lihir in its entirety would be treated as a single IU and all the clusters identified in this analysis would be subsumed into the same IU.

Little prior research has addressed the spatial-temporal epidemiology of yaws, and, to our knowledge, no prior research focuses on the spatial scale that is the focus of our analysis. A recent study from the Solomon Islands sampled villages and then households within villages to assess the prevalence of yaws and identify risk factors. The study found that yaws clusters in villages more so than in households but did not address spatial patterns at the scale of multiple neighboring villages [6]. Earlier work on yaws suggests that geographically heterogeneous environmental factors such as humidity influence the prevalence of yaws [25]. Environmental or social factors may lead to consistently higher levels of yaws in some locations: Public health staff on Lihir anecdotally described Tumbuapil—a village at the southern tip of Lihir as having a consistently high burden of yaws. Notably, Tumbuapil was part of statistically significant clusters in more active case surveys (4 of 7) than any other village and was part of the spatial-temporal cluster with the highest observed-to-expected ratio. We lack village-level data on risk factors, so we cannot assess what, if any, factors may contribute to higher prevalence in some villages rather than others.

Our study has several limitations: First, our results could reflect patterns in health-care seeking specific to yaws. While our analyses account for trends in the total number of outpatient visits from each village and condition on age and sex, spatially and temporally localized interest in seeking care for skin lesions could, in principle, generate such clusters but we think this is improbable. Second, an environmental or social risk factor for the transmissibility of yaws or people’s susceptibility to yaws could underpin the spatial-temporal clusters we observed. Multivillage spatial-temporal clusters in theory could form even in the absence of transmission between villages, but we are unaware of any risk factors that vary not only at the relevant spatial scale but also at the relevant temporal scale. Third, the number of outpatient visits varied greatly by village, with fewer visits from villages that are farther from LMC; we may have lacked power in our spatial-temporal analysis to detect yaws clusters in areas such as the northwest part of Lihir where we identified a spatial-only cluster in the seventh survey. Fourth, directly comparing the spatial-temporal and spatial-only clustering results is difficult in part because the analyses focus on somewhat different underlying populations: We had to include migrants in the spatial-only clustering analysis but could not include them in the spatial-temporal analysis. Additionally, our primary school attendance area clustering analysis associated each village with a single primary school and does not account for the fact that students from a single village may attend different primary schools. We used school attendance records from a single year to define primary school attendance areas; our analysis would not capture changing school attendance patterns. Finally, our findings may not be generalizable to other yaws-endemic areas. Elsewhere, villages may differ substantially from those on Lihir in terms of population size, density, proximity to other villages, and other factors. Conducting research on the spatial epidemiology of yaws in numerous settings would help identify phenomena that are contingent on a particular geography versus those that reflect general characteristics of yaws epidemiology.

Operational research in support of yaws eradication will need to continue to investigate the disease’s spatial epidemiology: Designing prevalence surveys that employ cluster sampling requires understanding the extent of spatial clustering of yaws. Moreover, a mechanistic understanding of the spatial and temporal scales at which yaws spreads and the extent to which transmission occurs in different settings (e.g. schools versus households) can help inform intervention strategies, particularly contact tracing. Earlier generations of yaws epidemiologists did not have access to molecular epidemiology tools or to remotely sensed data on environmental risk factors. These approaches may advance our understanding of the spatial epidemiology of yaws but will not replace the ongoing need for high-quality and high-spatial-resolution descriptive epidemiological data collected in both research and programmatic settings.

## Acknowledgements

We wish to thank Professor Marcia Castro for her advice on spatial statistics and Dr. Michael Marks for reviewing the manuscript and for helpful conversations on spatial implementation units and mapping strategy for yaws. We are grateful to the field, laboratory, and clinical staff at LMC for their efforts caring for patients and collecting data. Data on school attendance and geographic data on schools and villages were provided by the Sustainable Development office of Newcrest Mining Limited.

## Author Contributions

Conceptualization: EQM OM MBM.

Data Curation: EQM OM.

Formal analysis: EQM.

Funding acquisition: EQM OM.

Investigation: EQM OM.

Methodology: EQM.

Resources: OM.

Software: EQM.

Supervision: OM MBM.

Visualization: EQM

Writing - original draft: EQM.

Writing - review & editing: EQM OM MBM.

## Data Availability

The data and code necessary to replicate the analyses are available in the Harvard Dataverse at https://dataverse.harvard.edu/dataset.xhtml?persistentId=doi:10.7910/DVN/NZIVED.

## Supporting Information

**S1 Table.** Discrete Poisson analysis adjusted for age and sex, confirmed RPR-positive.

| ID | Start date | End date | Number of villages | Village IDs | Observed cases | Expected cases | p-value |
| --- | --- | --- | --- | --- | --- | --- | --- |
| 1 | 2010/8/2 | 2012/4/29 | 6 | Tumbuapil, Lissel, Komat, Lataul, Kinami, Pangoh | 97 | 21.56 | $<1 \times 10^{-17}$ |
| 2 | 2005/4/11 | 2009/3/1 | 4 | Putput_1, Putput_2, Lipuko, Matakues | 171 | 83.53 | $1.0 \times 10^{-11}$ |
| 3 | 2014/1/20 | 2016/5/29 | 4 | Kunaye_1, Kunaye_2, Kul, Zuen | 116 | 52.12 | $9.4 \times 10^{-9}$ |

**S2 Table.** Discrete Poisson analysis (unadjusted).

| ID | Start date | End date | Number of villages | Village IDs | Observed cases | Expected cases | p-value |
| --- | --- | --- | --- | --- | --- | --- | --- |
| 1 | 2010/8/2 | 2012/3/11 | 6 | Tumbuapil, Lissel, Komat, Lataul, Kinami, Pangoh | 104 | 28.90 | $<1 \times 10^{-17}$ |
| 2 | 2005/4/11 | 2008/6/8 | 4 | Lipuko, Putput_2, Matakues, Putput_1 | 178 | 79.79 | $1.2 \times 10^{-14}$ |
| 3 | 2009/8/17 | 2014/12/14 | 2 | Kunaye_1, Kunaye_2 | 214 | 119.75 | $2.7 \times 10^{-9}$ |

**S3 Table.** Space-time permutation analysis adjusted for age and sex.

| ID | Start date | End date | Number of villages | Village IDs | Observed cases | Expected cases | p-value |
| --- | --- | --- | --- | --- | --- | --- | --- |
| 1 | 2010/8/2 | 2012/3/11 | 6 | Tumbuapil, Lissel, Komat, Lataul, Kinami, Pangoh | 104 | 44.14 | $9.0 \times 10^{-13}$ |
| 2 | 2005/4/11 | 2008/6/1 | 4 | Lipuko, Putput_2, Matakues, Putput_1 | 177 | 103.80 | $5.3 \times 10^{-9}$ |
| 3 | 2012/5/7 | 2016/5/29 | 5 | Kul, Kunaye_1, Kunaye_2, Londolovit, Zuen | 240 | 159.64 | $2.5 \times 10^{-7}$ |

**S4 Table.** Space-time permutation analysis (unadjusted).

| ID | Start date | End date | Number of villages | Village IDs | Observed cases | Expected cases | p-value |
| --- | --- | --- | --- | --- | --- | --- | --- |
| 1 | 2010/8/2 | 2012/3/11 | 6 | Tumbuapil, Lissel, Komat, Lataul, Kinami, Pangoh | 104 | 43.21 | $3.0 \times 10^{-12}$ |
| 2 | 2005/4/11 | 2008/6/1 | 4 | Lipuko, Putput_2, Matakues, Putput_1 | 177 | 102.24 | $9.4 \times 10^{-9}$ |
| 3 | 2012/5/7 | 2016/5/29 | 5 | Kul, Kunaye_1, Kunaye_2, Londolovit, Zuen | 240 | 157.44 | $3.0 \times 10^{-7}$ |

**S5 Table.** P-values for primary school attendance area analysis. These p-values result from the permutation test used to assess clustering of yaws in 8- to 14-year-olds by primary school attendance area for each year from 2006 through 2015 and for the entire study period.

| **year** | **p-value** |
| --- | --- |
| 2006 | 0.0687 |
| 2007 | 0.4778 |
| 2008 | 0.4068 |
| 2009 | 0.9881 |
| 2010 | 0.6705 |
| 2011 | 0.5949 |
| 2012 | 0.9315 |
| 2013 | 0.9808 |
| 2014 | 0.6875 |
| 2015 | 0.9795 |
| entire study period | 0.6846 |

**S1 Fig.**
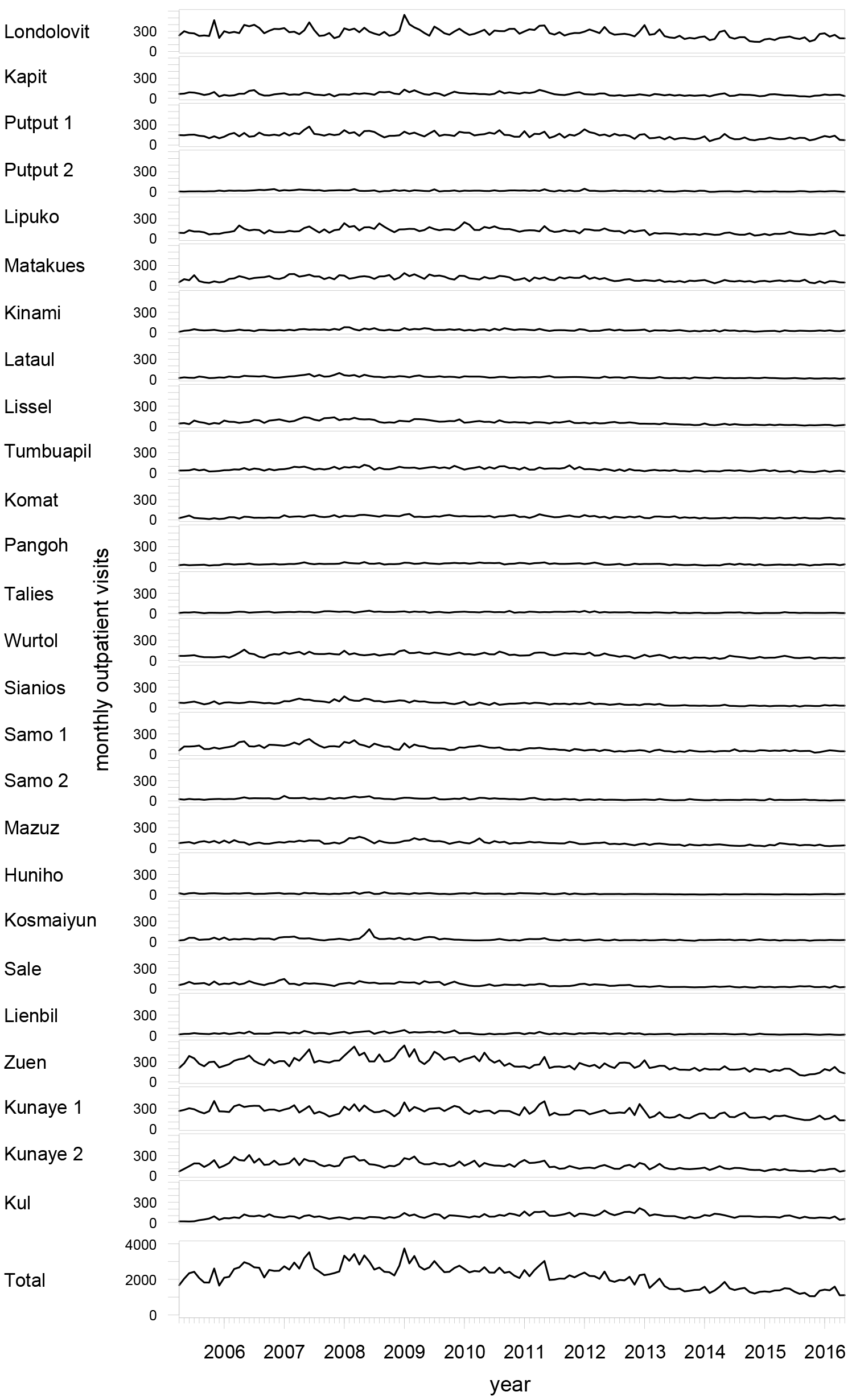
Outpatient visits to LMC. Time series of outpatient visits (any diagnosis) aggregated by month by village and for all villages combined. The villages are ordered to match their sequential order around the circumference of Lihir.

**S2 Fig.**
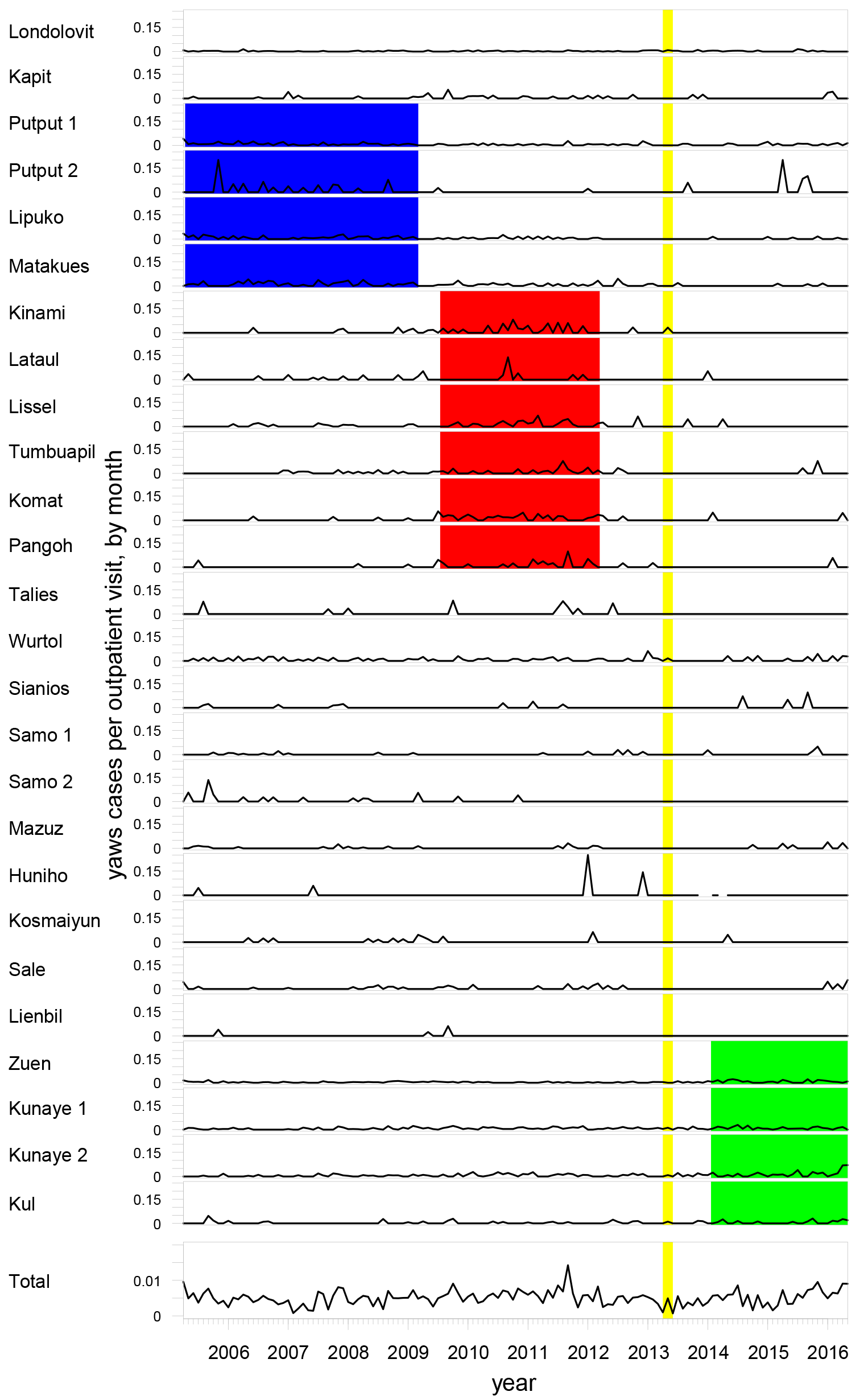
Passively detected yaws cases as a proportion of outpatient visits. The time series show the proportion of outpatient yaws diagnoses as a fraction of all outpatient visits at LMC aggregated by month by village and for all villages combined. The villages are ordered to match their sequential order around the circumference of Lihir. Red, blue, and green rectangles correspond to spatial-temporal clusters 1, 2, and 3, respectively, from Fig 3. The vertical yellow bar corresponds to when mass drug administration was implemented on Lihir.

**S3 Fig.**
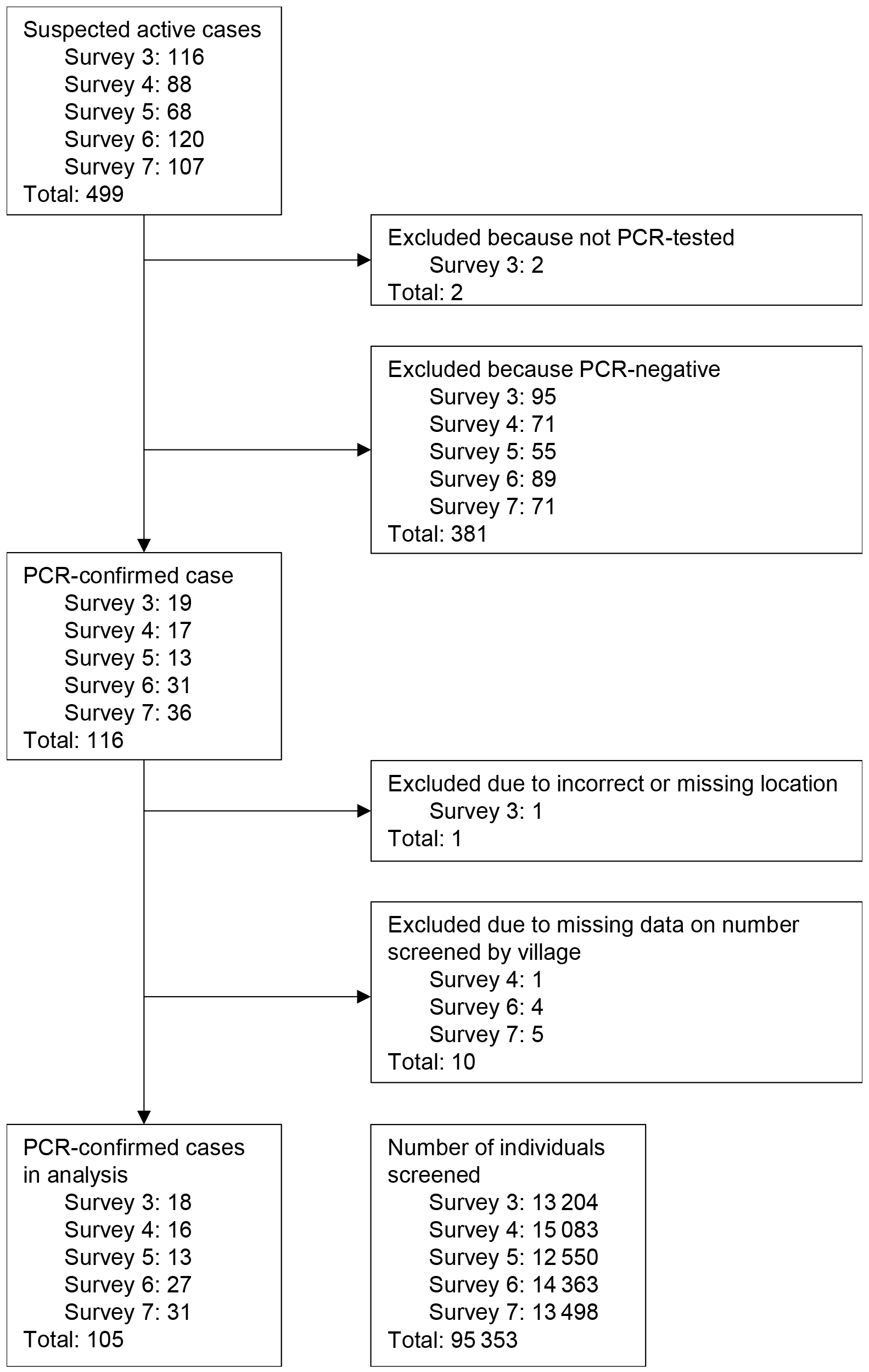
Flow diagram for data in analysis of PCR-confirmed prevalent yaws cases. The diagram lists by active case finding survey the number of suspected active cases, PCR-confirmed cases, excluded cases, PCR-confirmed cases in the final analysis, and number of individuals screened.

**S4 Fig.**
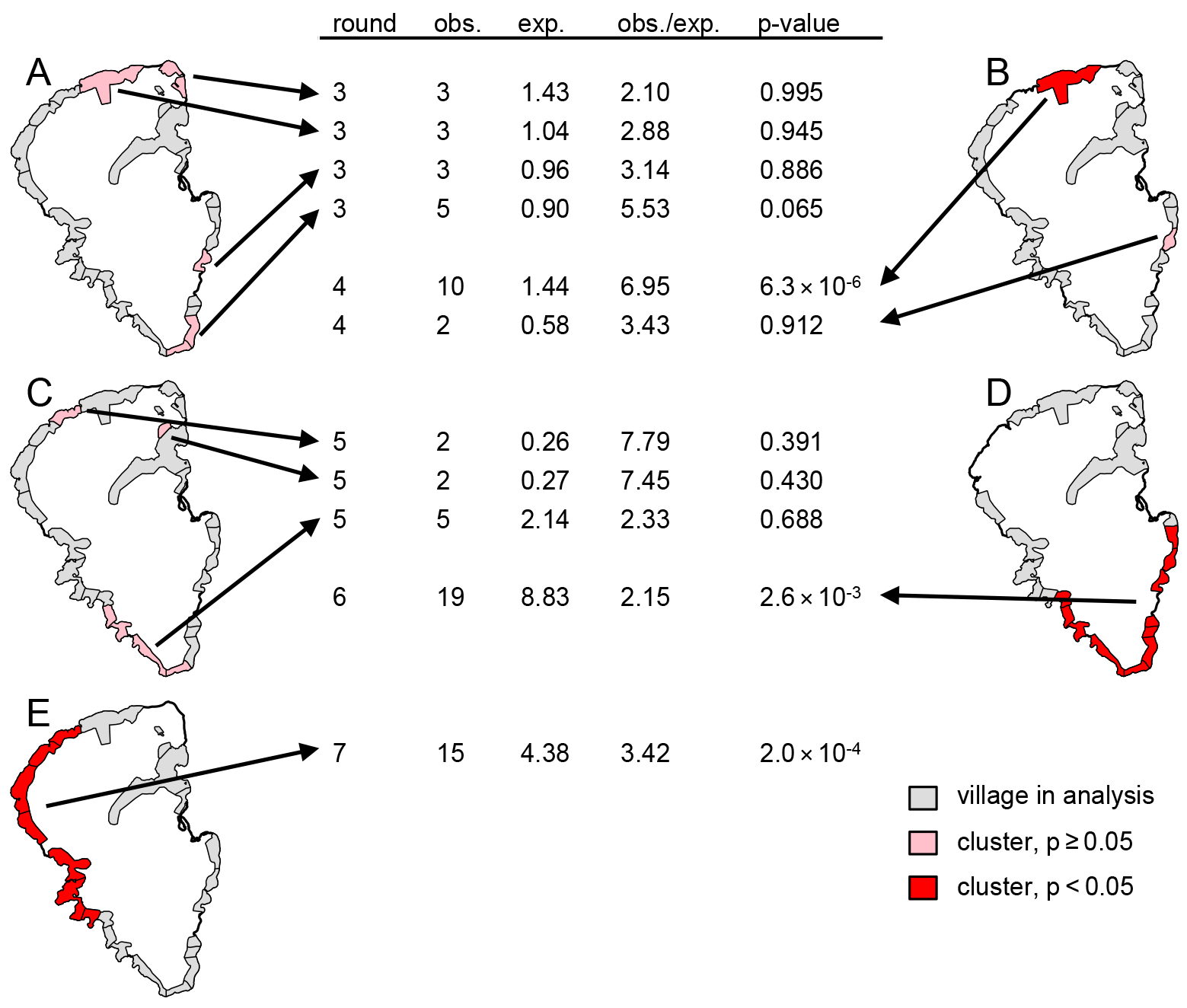
Spatial clusters of actively detected PCR-confirmed yaws cases. Results of spatial-only discrete Poisson SaTScan analysis for PCR-confirmed prevalent yaws cases identified via active case finding for each survey 3 through 7 (A ‒ E, respectively). Villages that are part of statistically significant spatial clusters are shaded in red and not statistically significant clusters are shaded in pink. Villages are shaded gray if they were part of the analysis in the corresponding survey (but not identified as part of a cluster). Villages are excluded from the map for each survey where the number of individuals screened in that survey in that village is unknown. The arrows point from each cluster to details describing the number of observed yaws cases in that cluster, the number of expected yaws cases, the ratio of observed to expected, and p-value for that cluster.

## References

1. Mitjà O, Asiedu K, Mabey D. Yaws. Lancet. 2013;381(9868):763–73.

2. Mitjà O, Marks M, Konan DJP, Ayelo G, Gonzalez-Beiras C, Boua B, et al. Global epidemiology of yaws: a systematic review. Lancet Glob Heal. 2015;3(6):e324–31.

3. Mitjà O, Hays R, Ipai A, Penias M, Paru R, Fagaho D, et al. Single-dose azithromycin versus benzathine benzylpenicillin for treatment of yaws in children in Papua New Guinea: an open-label, non-inferiority, randomised trial. Lancet. 2012;379(9813):342–7.

4. World Health Organization. Eradication of yaws - the Morges Strategy. Wkly Epidemiol Rec. 2012;87(20):189–94.

5. World Health Organization. Global Programme To Eliminate Lymphatic Filariasis Monitoring and Epidemiological Assessment of Mass Drug Administration. 2011.

6. Marks M, Vahi V, Sokana O, Puiahi E, Pavluck A, Zhang Z, et al. Mapping the epidemiology of yaws in the Solomon Islands: a cluster randomized survey. Am J Trop Med Hyg. 2015;92(1):129–33.

7. Mitjà O, Hays R, Ipai A, Gubaila D, Lelngei F, Kirara M, et al. Outcome predictors in treatment of yaws. Emerg Infect Dis. 2011;17(6):1083–5.

8. Marks M, Mitjà O, Vestergaard LS, Pillay A, Knauf S, Chen CY, et al. Challenges and key research questions for yaws eradication. Lancet Infect Dis. 2015;15(10):1220–5.

9. Bejon P, Williams TN, Nyundo C, Hay SI, Benz D, Gething PW, et al. A micro-epidemiological analysis of febrile malaria in coastal Kenya showing hotspots within hotspots. Elife. 2014;2014(3):1–13.

10. Koch R. Zusammenfassende Darstellung der Ergebnisse der Malariaexpedition. Dtsch Medizinische Wochenschrift. 1900;26(49):781–3.

11. Bainton NA. The Lihir Destiny: Cultural Responses to Mining in Melanesia. Canberra: ANU E Press; 2010.

12. U.S. Geological Survey, National Geospatial-Intelligence Agency, National Aeronautics and Space Administration. Shuttle Radar Topography Mission 1 Arc-Second Global: SRTM1S04E152V3. Sioux Falls, South Dakota: U.S. Geological Survey (USGS) Earth Resources Observation and Science (EROS) Center; 2014.

13. GADM. Global Administrative Areas v. 2.0 [Internet]. 2011. Available from: http://www.gadm.org/

14. Marks M, Mitjà O, Solomon AW, Asiedu KB, Mabey DC. Yaws. Br Med Bull. 2015;113(1):91–100.

15. Kulldorff M. A spatial scan statistic. Commun Stat - Theory Methods. 1997;26(6):1481–96.

16. Kulldorff M, Information Management Services. SaTScanTM v9.4: Software for the spatial and space-time scan statistics. [Internet]. 2016. Available from: http://www.satscan.org/

17. Kleinman K. rsatscan: Tools, Classes, and Methods for Interfacing with SaTScan StandAlone Software [Internet]. 2015. Available from: https://cran.r-project.org/package=rsatscan

18. Mitjà O, Houinei W, Moses P, Kapa A, Paru R, Hays R, et al. Mass treatment with singledose azithromycin for yaws. N Engl J Med. 2015;372(8):703–10.

19. Mitjà O, Lukehart SA, Pokowas G, Moses P, Kapa A, Godornes C, et al. Haemophilus ducreyi as a cause of skin ulcers in children from a yaws-endemic area of Papua New Guinea: A prospective cohort study. Lancet Glob Heal. 2014;2(4):235–41.

20. Agresti A, Coull BA. Approximate is better than “exact” for interval estimation of binomial proportions. Am Stat. 1998;52(2):119–26.

21. Chasalow S. combinat: Combinatorics Utilities [Internet]. 2012. Available from: https://cran.r-project.org/package=combinat

22. R Core Team. R: A Language and Environment for Statistical Computing [Internet]. Vienna, Austria; 2014. Available from: http://www.r-project.org/

23. Kulldorff M, Heffernan R, Hartman J, Assunção R, Mostashari F. A space-time permutation scan statistic for disease outbreak detection. PLoS Med. 2005;2(3):0216–24.

24. Mitjà O, Godornes C, Houinei W, Kapa A, Paru R, Abel H, et al. Re-emergence of yaws after single mass azithromycin treatment followed by targeted treatment: A longitudinal study. Lancet. 2018;6736(18):1–9.

25. Hill KR. Non-specific factors in the epidemiology of yaws. Bull World Heal Organ. 1953;8(1-3):17–51.

